# Chromatin Mechanics Dictates Subdiffusion and Coarsening Dynamics of Embedded Condensates

**DOI:** 10.1101/2020.06.03.128561

**Authors:** Daniel S.W. Lee, Ned S. Wingreen, Clifford P. Brangwynne

**Affiliations:** Lewis-Sigler Institute for Integrative Genomics, Princeton University, Princeton, New Jersey 08544, USA; Department of Chemical and Biological Engineering, Princeton University; Department of Molecular Biology, Princeton University; The Howard Hughes Medical Institute

## Abstract

DNA is organized into chromatin, a complex polymeric material which stores information and controls gene expression. An emerging mechanism for biological organization, particularly within the crowded nucleus, is biomolecular phase separation into condensed droplets of protein and nucleic acids. However, the way in which chromatin impacts the dynamics of phase separation and condensate formation is poorly understood. Here, we utilize a powerful optogenetic strategy to examine the interplay of droplet coarsening with the surrounding viscoelastic chromatin network. We demonstrate that droplet growth dynamics are directly inhibited by the chromatin-dense environment, which gives rise to an anomalously slow coarsening exponent, *β*∼0.12, contrasting with the classical prediction of *β*∼ 1/3. Using scaling arguments and simulations, we show how this arrested growth can arise due to subdiffusion of individual condensates, predicting *β*∼α/3, where α is the diffusion exponent. Tracking the fluctuating motion of condensates within chromatin reveals a subdiffusive exponent, α∼0.5, which explains the anomalous coarsening behavior and is also consistent with Rouse-like dynamics arising from the entangled chromatin. Our findings have implications for the biophysical regulation of the size and shape of biomolecular condensates, and suggest that condensate emulsions can be used to probe the viscoelastic mechanical environment within living cells.

## Introduction

Liquid-liquid phase separation (LLPS) has emerged as a common assembly mechanism within living cells^1,2^. LLPS drives formation of a multitude of biomolecular condensates such as nucleoli^3^, stress granules^4^, and P granules^5^. Phase separation of proteins typically relies on multivalent interactions, and many efforts have been made to understand the role of self-interactions via intrinsically disordered protein regions (IDRs)^6^, and heterotypic interactions, often mediated by folded protein-protein interaction domains and oligomerized RNA (or other substrate) binding domains. The condensates that result from these interactions are multicomponent structures, which range from compositionally homogeneous liquid-like structures, to multiphase liquids and kinetically arrested gel states^7–9^. Their structural features and material properties are thought to enable a diversity of functions, ranging from increasing reaction rates by concentrating reactants, to inhibiting reactivity through sequestration, and even structurally rearranging their environment^1,10^.

Many phase-separated condensates appear to exhibit stable sizes dependent on cell type^11,12^. It is unclear how these sizes are maintained, since liquid compartments should not have an intrinsic size scale. Rather, given enough time, individual liquid condensates should coalesce to form a single, large droplet. In general, following a full quench of the system, i.e., after nucleation and depletion of the dilute phase, the mean droplet radius is expected to evolve as a power law with exponent *β*, such that ⟨*R*⟩∼*t*^β^ where *β* = 1/3. Two distinct processes identically predict this exponent: Lifshitz-Slyozov Wagner (LSW) theory describes growth by Ostwald ripening^13^, and Brownian-motion driven coalescence (BMC) describes how individual droplets collide and fuse^14^. Droplet growth obeying this 1/3 power law has been observed in a variety of biological systems, including lipid membranes^15^, *in vitro* protein droplets, and nucleoli in *C. elegans* embryos^16^, but distinguishing between the two mechanisms is not always possible.

In addition to the ambiguity in distinguishing these two processes, our understanding of the physics governing condensate coarsening within living cells is further complicated by the strongly viscoelastic nature of the intracellular environment. Indeed, droplet movements and interactions are expected to be strongly impacted by their viscoelastic environment. A particularly striking example of this interplay is found in *Xenopus laevis* oocytes, whose nuclei contain hundreds of nucleoli, which are strongly constrained by an F-actin scaffold that only relaxes on very long time scales, giving rise to a broad power-law distribution of nucleolar sizes^3^. Upon actin disruption, the nucleoli sediment under gravity and undergo large scale coalescence into a single massive nucleolus^17^. In human cell lines, nucleoli exhibit liquid-like properties of formation and fusion but vary in their shapes and sizes^18^. Interestingly, these size and shape variations are tied to nuclear processes such as transcription and chromatin condensation^19^, suggesting that active processes can influence the properties and distribution of phase separated condensates. Taken together, these findings are consistent with the shapes and sizes of nuclear condensates being strongly influenced by their physical environment.

In addition to nucleoli, the nucleus contains dozens of different types of condensates, including Cajal bodies, paraspeckles, PML bodies, and nuclear speckles, each of which is intimately associated with the surrounding chromatin network. Chromatin is the active polymeric material into which DNA and its associated proteins assemble and is packed into the nucleus at dense volume fractions, with measurements ranging from 12 to 52%^20^. Chromatin can be considered a continuous material on the length scale of microscopically visible droplets: electron microscopy images^20^, as well as experiments utilizing microinjected fluorescent dextrans^21^, both suggest an average chromatin pore size of less than 100 nm^22^, although chromatin exhibits significant heterogeneity on micron length scales^23^. Micromanipulation of extracted nuclei has also revealed that chromatin exhibits its own force response independent of the nuclear lamina, which has been the focus of most nuclear mechanics studies. The mechanical response of chromatin appears to depend on its epigenetic modifications and degree of compaction^24^, although chromatin’s frequency-dependent viscoelastic response has not been studied in detail. Despite the clear functional importance of understanding the link between the dynamics of phase separation and the surrounding viscoelastic genetic material, very few studies have examined this connection.

Most studies to date have relied on simplified *in vitro* systems using purified components in an aqueous buffer. As a result, the impact of the cellular milieu on phase separation has generally been unexplored, partly due to a lack of tools for interrogating phase separation in living cells^25^. However, recent advances have led to the development of optogenetic approaches, which utilize light to control intracellular phase separation through oligomerization of disordered and substrate binding domains^7,22,26,27^. Experiments using these systems have suggested that nuclear condensates preferentially form in relatively chromatin-poor regions and tend to exclude chromatin as they grow^22^. However, the physics governing the relationship between chromatin’s material properties, and the formation, mobility, growth, and ultimate size distribution of embedded condensates, remains unclear.

Here, we utilize the biomimetic Corelet system^27^ to generate model IDR-based condensates and examine their coarsening dynamics and interactions with chromatin in living cells. We demonstrate that coarsening in the nucleus is significantly slower than classical theory predicts, and occurs primarily via coalescence, rather than via Ostwald ripening. Using particle tracking experiments, we show that this slow coarsening results from strongly subdiffusive condensate motion, in quantitative agreement with a theoretical prediction relating the coarsening and diffusive exponents. The growth dynamics of embedded condensates are thus sensitively dependent on the viscoelasticity of the surrounding chromatin, representing a novel emulsion-based readout of the mechanical environment within living cells.

## Results

### Optogenetically engineered condensates exhibit slow coarsening dynamics

To study interactions between proteinaceous condensates and chromatin, we utilized the two-component Corelet system^27^ (Fig. 1), to control condensate formation in living cells. Corelets are based on a ferritin core composed of 24 subunits each fused to an iLiD domain, and a phase separation-driving IDR fused to sspB, iLiD’s optogenetic heterodimerization partner. Here, we used the IDR from the Fused in Sarcoma (FUS) protein, a member of the FET family of transcription factors^28^. We expressed these constructs in human osteosarcoma (U2OS) cells, which contain nuclei typical of mammalian cells, and yet are experimentally tractable, particularly for microscopy studies. Consistent with previous work^27^, upon blue light activation, iLiD and sspB bind, resulting in oligomerized IDR assemblies that phase separate above a threshold of concentration and valence (degree of oligomerization). These model phase-separated nuclear condensates are visible within seconds of blue light activation, providing a powerful platform for spatially and temporally interrogating the impact of the viscoelastic environment on their dynamic coarsening behavior.

**Fig. 1:**
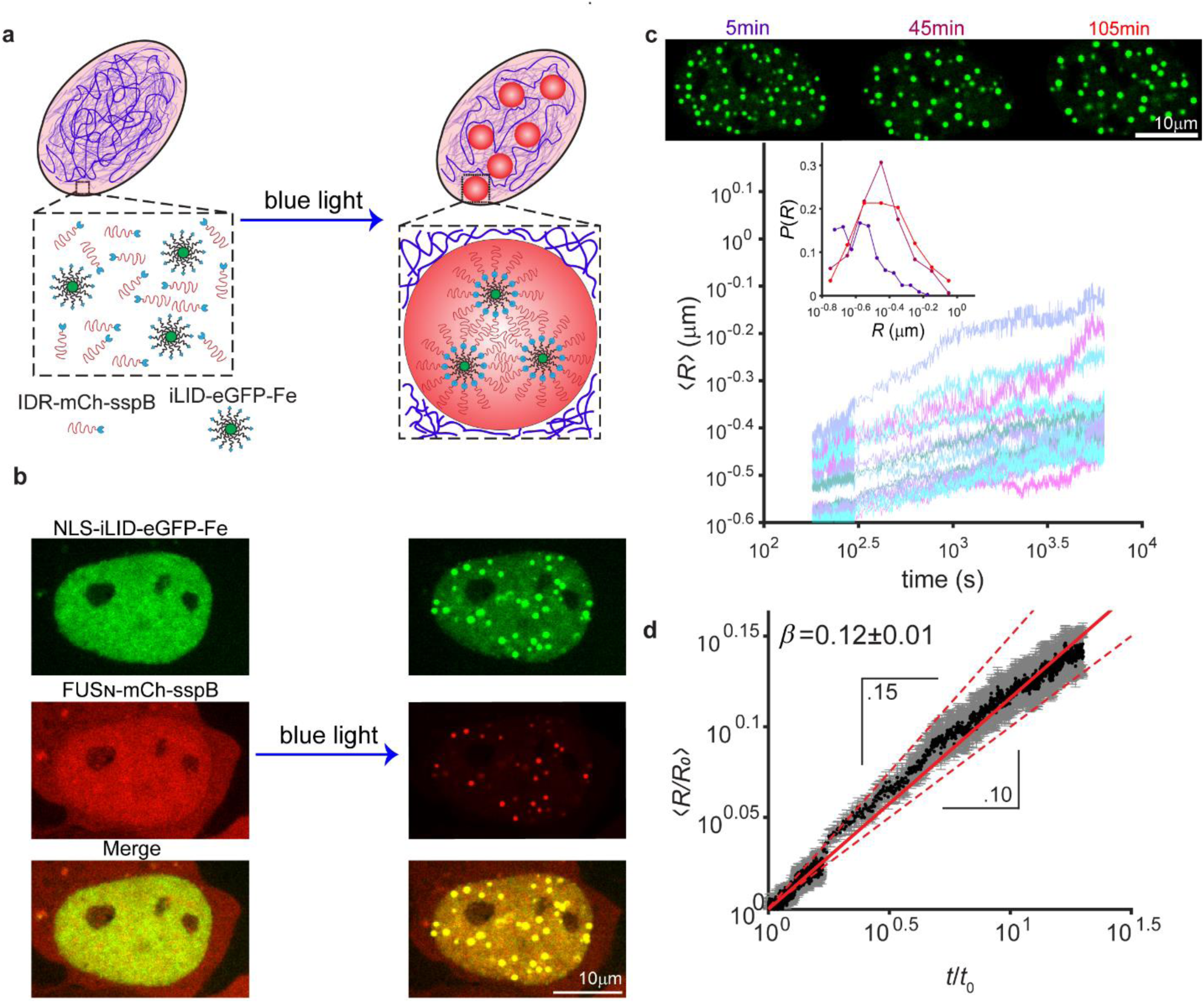
Coarsening dynamics of light-activated protein condensates in living cells. **a,** In the presence of blue light, the Corelet system recruits IDRs (red) to the ferritin cores (green), to drive phase separation in the nucleus of living cells. **b**, Within seconds of exposure to blue light, droplets nucleate and grow. **c,** Top images show snapshots of FUS Corelet condensates at indicated times. Main plot shows average droplet radii for 18 cells calculated over 105 minutes and plotted starting 3 minutes after activation. The size distribution for 8 cells with similar supersaturation (mean supersaturation of 0.77 with standard deviation 0.08) was taken at timepoints of 5, 45, and 105 minutes. **d,** The average radius of droplets per cell ⟨*R*/*R*_0_⟩ was plotted over time, after normalizing by the radius *R*_0_ of each droplet at the defined *t*_0_ of 3 minutes, revealing a power law with exponent approximately 0.12 (solid red line). Error was propagated as standard error of the mean (SEM). Dashed red lines have slopes of 0.15 and 0.10, respectively.

We first sought to characterize droplet coarsening in live nuclei, by activating cells and measuring Corelet droplets over the course of 105 minutes. At very early times, coarsening could be convoluted with nucleation and growth, initially driven by the supersaturated dilute phase, with an early-stage growth exponent predicted to be 1/2^29^. We found that the dilute-phase intensity rapidly decreased and began stabilizing after roughly 3 minutes of activation; a slow subsequent linear decrease is likely due to photobleaching (Supplementary Fig. 2). We thus wait until after this transient dilute-phase equilibration and then probe the coarsening dynamics by determining the mean radius of droplets over time (Fig. 1d). Averaging over cells, we find a consistent power-law exponent of *β* = 0.12 ± 0.01(Fig. 1e). This low exponent is surprising, given that both Ostwald ripening and Brownian motion coalescence-dominated coarsening predict *β* = 0.33, which is commonly observed in nonliving systems^15^. Our results are not an artifact of the small size of droplets, as similar results were found by using integrated intensity as a proxy for size (Supplementary Fig. 1). Moreover, the observed heterogeneity across cells in initial droplet sizes is accounted for by variation in the expression of protein constituents; individual fits on a per-cell basis show that scaling behavior follows a similar exponent, with more noise at low volume fraction (Supplementary Fig. 2).

**Fig. 2:**
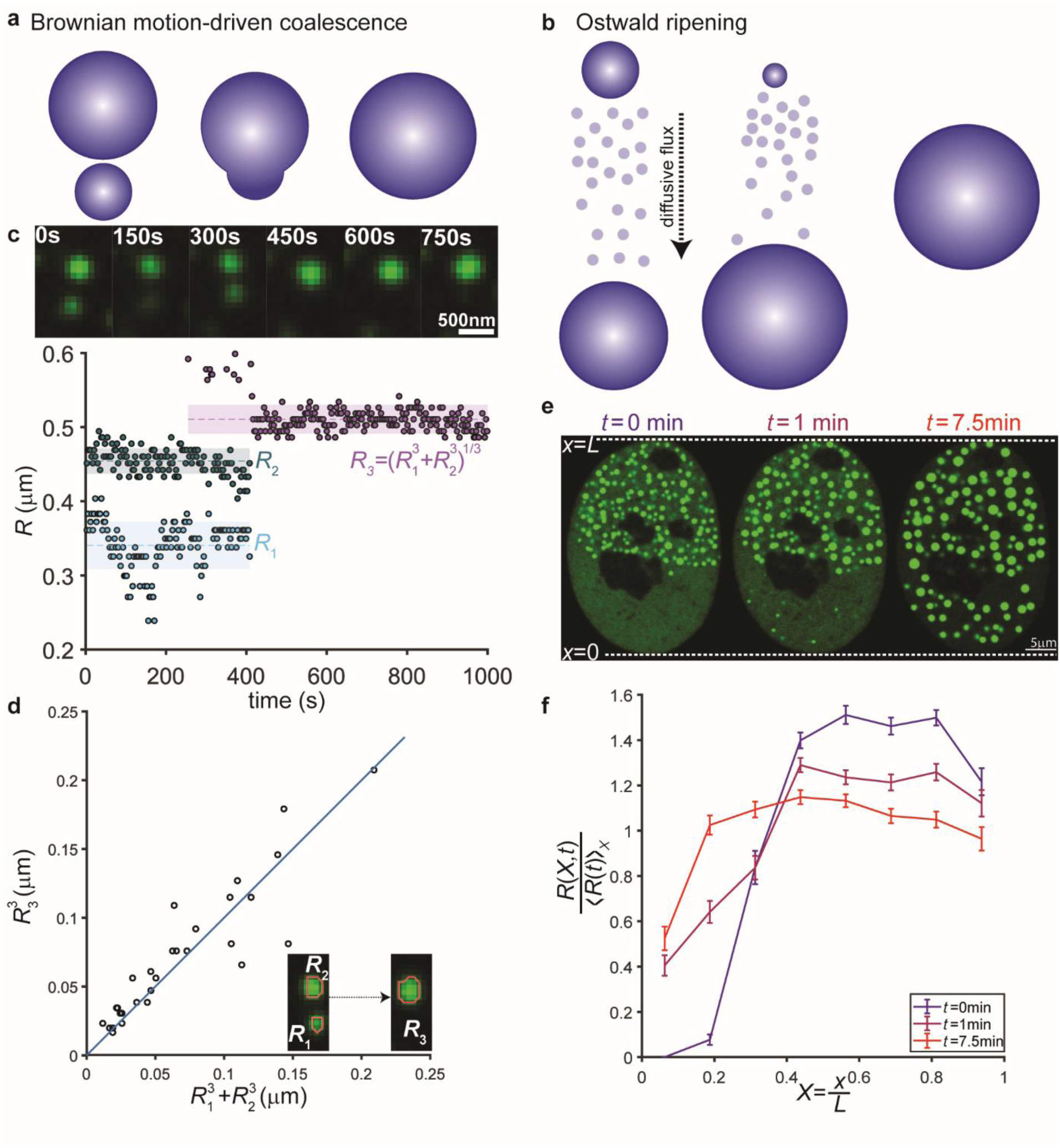
Droplet growth is dominated by coalescence. **a,b,** Emulsions generally coarsen by Ostwald ripening or by mergers, respectively characterized by continuous and discrete growth. **c,** Example (plot with select corresponding images) showing condensates are stable over long timescales, remaining the same size for minutes and then coming into contact (cyan and green points), merging, and settling at a stable size consistent with volume conservation (black line shows predicted conserved volume of final droplet; uncertainty of prediction is propagated from standard errors of the time-averaged means of the original two droplets). **d,** Volume conservation accounts for size changes over many collisions (n=29). **e,** Gradient activations were used to generate polydisperse droplets. Following 30 seconds of activation with an intensity gradient of blue light, droplets formed primarily on the high intensity side of the nucleus. After switching to global activation, new smaller droplets nucleated and grew on the other side of the nucleus. **f,** Quantification of the experiment described in **e** over 10 cells. Droplets were segmented and binned by location along the axis of the nucleus; droplet size was normalized for each cell and averaged over cells. Error bars are SEM.

### Droplet coarsening is dominated by coalescence events

To elucidate the mechanism underpinning this anomalously slow growth, we investigated the relevance of two possible scenarios of droplet coarsening, Ostwald ripening (i.e. LSW) and Brownian-motion-driven coalescence (BMC) (Fig. 2a,b). While both mechanisms are driven by the minimization of surface energy, Ostwald ripening would occur via continuous growth/shrinkage of droplets, while a merger-dominated process would take place via discrete jumps in size associated with collision events. Consequently, we tracked pairs of droplets that merged, finding that their sizes were stable for minutes before merger, at which point the resulting merged droplet attained the size predicted by volume conservation and remained that size for minutes in the absence of further collisions (Fig. 2c). This behavior was consistent over 29 apparent mergers analyzed across 18 cells, based on single bookending frames independently segmented before and after identified in-plane merger events (Fig. 2d), suggesting that growth over the entire bookended time interval (648±62 seconds on average) was accounted for by coalescence.

These data are consistent with BMC-dominated coarsening, albeit with an anomalously low exponent. However, we reasoned that Ostwald ripening could also play a role, but would manifest relatively infrequently, in shrinkage and growth of unequal-sized droplets. To create a particularly polydisperse distribution, we activated nuclei with a low-intensity spatial gradient of blue light for 30 seconds, resulting in nuclei with droplets accumulated on one side and a local enrichment of protein in the dilute phase on the activated side, consistent with previously observed ‘diffusive capture’ of core molecules^9^. Switching to uniform blue light stimulation resulted in fast equilibration of the dilute phase (<10 seconds, Supplementary Fig. 3) and therefore a uniform influx of newly-activated molecules. Within minutes, we observed nucleation and growth of droplets on the previously unactivated side (Fig. 2e,f). We reasoned that if Ostwald ripening played an important role in the growth of our droplets, any small, newly nucleated droplets would quickly vanish since ripening would favor the preexisting ones, which are on the order of 3-5 fold larger in radius. However, we observed the opposite, finding that the new droplets grew to a similar size as the preexisting ones. Thus, we conclude that late-stage growth of our droplets is primarily driven by droplet coalescence events.

**Fig. 3:**
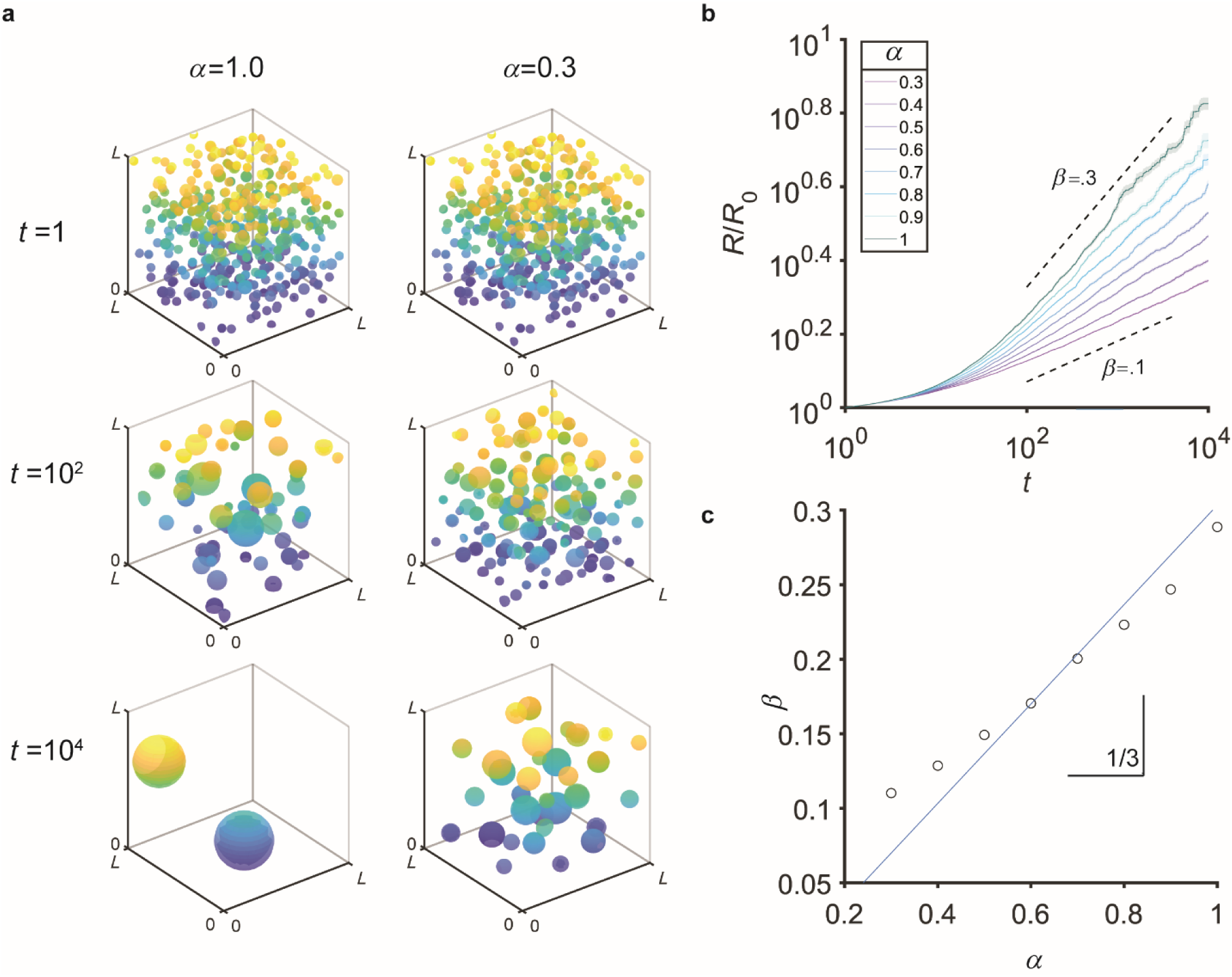
Simulations quantitatively demonstrate the scaling relation between the diffusive exponent and the coarsening exponent. **a,** 500 droplets were placed in a three dimensional box at a volume fraction of 0.05 and allowed to diffuse with a specified subdiffusive exponent *α*, and merge upon collision. **b,** Values of *α* were chosen ranging from 0.3 to 1; for each of twenty replicates per condition, the average radius of the droplets was calculated, normalized to the average at *t* = 0. Shaded region indicates standard error of the mean. **c,** For each simulation condition, the coarsening exponent was calculated by fitting a power law starting at *t* = 10^2^ and plotted against the input diffusive exponent *α*.

### Coarsening exponent is predicted to depend on diffusive exponent

Having empirically concluded that the coarsening of droplets is primarily driven by coalescence, we sought to quantitively explain the anomalously slow coarsening exponent. We therefore revisited scaling arguments from the literature^15^ which assume ordinary diffusion of droplets. More generally, the rate at which droplets collide should depend on their mean squared displacement (MSD), i.e. ⟨*x*^2^⟩∼*D*_*r*_Δ*t*^α^ where *α* is the diffusion exponent (*α* = 1 for ordinary diffusion, *α* < 1 for subdiffusive motion, e.g. in a viscoelastic environment). At fixed droplet volume fraction (i.e., having reached a steady-state dilute-phase concentration), the typical spacing *l* between droplets is related to their typical radius *r* by *l*∼*r*. Then, assuming that droplets collide over a time scale at which their mean-squared displacement equals their typical spacing, i.e. ⟨*x*^2^⟩∼*l*^2^, and assuming a diffusion coefficient *D*_*r*_∼*r*^−1^ as in Stokes drag, we find that *r*(*t*)∼*t*^α/3^, i.e., *β* = α/3, confirming our intuition that subdiffusing droplets should coarsen slowly. Note that, in the case of ordinary diffusion (α = 1), we recover the expected *β* = 1/3.

To examine the validity of this scaling argument, we performed simulations on the coarsening of subdiffusing droplets. We generated 500 droplets of identical initial radius, randomly placed in a 3D continuous space with periodic boundary conditions at a volume fraction of 5%. Droplets diffused according to fractional Brownian motion (fBM), with a step size inversely proportional to their radius such that *D*_*r*_∼*r*^−1^. After initial placement and subsequently after each timestep, overlapping droplets were merged with volume conserved and the new droplet placed at their center of mass. Unsurprisingly, the more subdiffusive the motion, the slower the coarsening of the droplet size distribution (Fig. 3a). The average droplet radius over time was calculated for each simulation (Fig. 3b), and the coarsening exponent *β* was obtained by fitting a power law starting at *t*=10^2^. We found that *β* ≈ 0.3α, in good agreement with our scaling argument.

### Droplets exhibit subdiffusive motion due to constraints from surrounding chromatin

Our simulations and scaling argument suggest that the surprisingly small coarsening exponent *β* could arise from subdiffusive droplet dynamics. Since the nucleus contains a dense chromatin network, we reasoned that droplets may indeed exhibit subdiffusive dynamics due to the presence of chromatin. We therefore first examined the physical interactions between the droplets and chromatin. To generate droplets of a size significantly above the diffraction limit we applied a local light activation protocol. We illuminated an approximately 2-micron diameter circular region for 10 minutes, in cells stained with Hoechst 33342, a marker of bulk DNA density. Consistent with previous experiments using the related CasDrop system, we observed depletion of the DNA marker from the droplet, suggesting that chromatin is largely excluded from the droplets (Fig. 4a,b). We found that IDRs from several transcriptionally relevant proteins, including those from HNRNPA, TAF15, and BRD4, show a similar chromatin-exclusion effect (Fig. 4c). Qualitatively, all these large droplets moved relatively little, over minutes of observation. Likewise, following global activation for well over an hour, smaller droplets remained distributed throughout the nucleus, with their spatial position changing very little (Fig. 1c). We therefore surmised that droplets are constrained by the surrounding chromatin network.

**Fig. 4:**
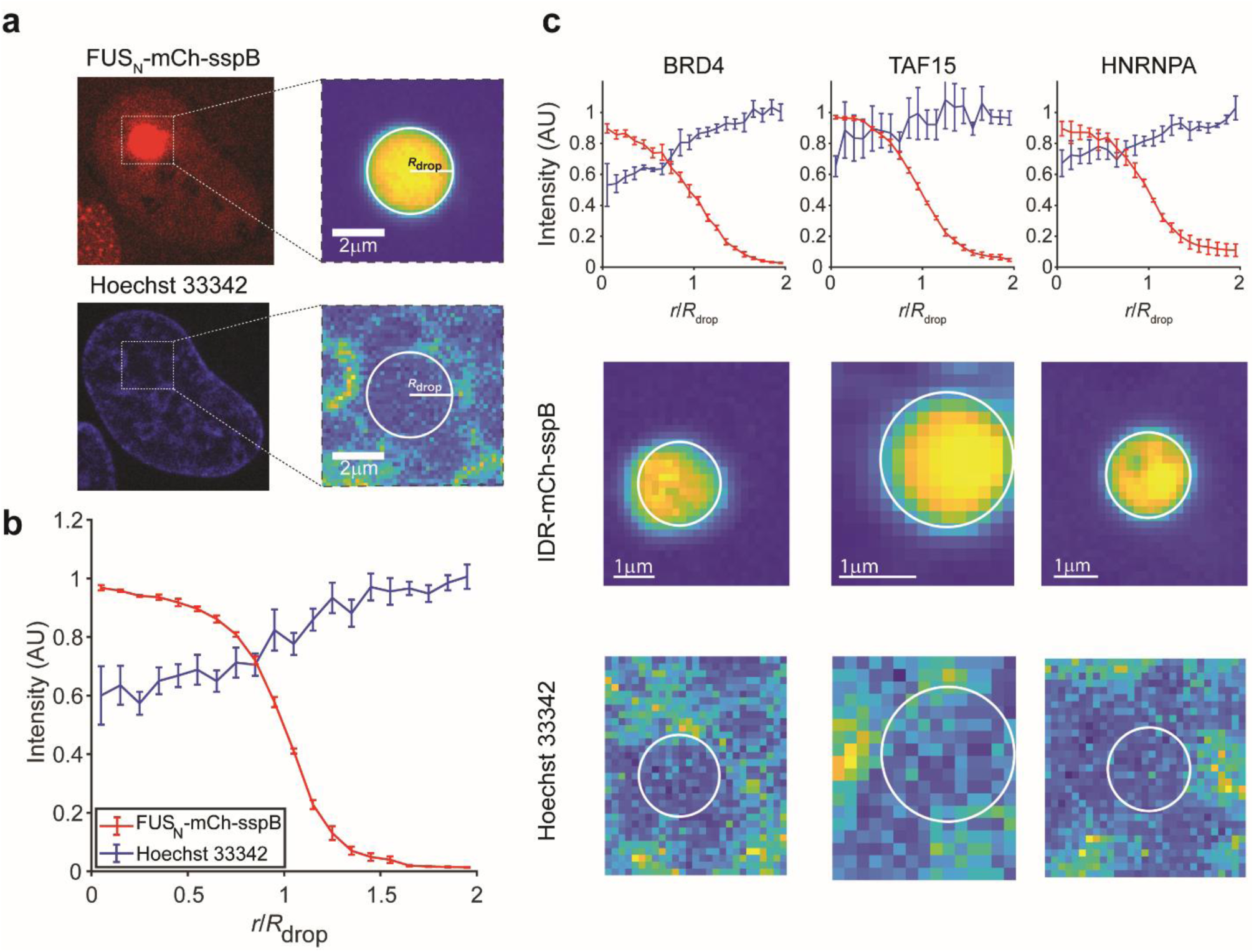
Droplets occupy chromatin-poor regions. **a,** Droplets can be locally ‘written’ in specific locations by patterning blue light stimulation. Co-staining with Hoechst 33342 demonstrates that droplets are generally associated with chromatin-sparse regions. **b,** Radially averaged intensity profiles of the Hoechst and droplet channels demonstrate that the chromatin is depleted within the droplet (averaged over 5 droplets). **c**, Identical analyses on droplets formed using three other IDRs (BRD4dN, TAF15_N,_ HNRNPA_C_) reveal similar exclusion of chromatin. Error bars show SEM over 5, 7, and 5 droplets respectively.

To quantitatively test the predicted relationship between the coarsening and diffusive exponents, i.e. *β* = α/3, we performed particle tracking on individual droplets to characterize their motion. Droplets were tracked over two timescales, at a rate of 60 ms/frame for 2 minutes and later at a rate of 3 seconds/frame for 100 minutes. The plotted MSDs were calculated by first time averaging within trajectories and then ensemble averaging over all droplets for multiple cells. Over long timescales (several minutes), we found that bulk translation of the nucleus appeared to increase the MSD; the diffusive exponent was therefore taken from the early times. We find that droplets do, in fact, exhibit subdiffusive motion: from a fit over lag times Δ*t* ranging from 60ms to 12s, we obtain an exponent of approximately α = 0.48 ± .08.

This anomalous exponent α was calculated using ensemble-averaged MSDs taken from droplets generated with a global activation protocol and relatively low variation in size. Since α could depend on size, for example if droplet size were comparable to the chromatin mesh size, we sought to examine this aspect further. We applied local activation as described above to generate droplets of greatly varying sizes (ranging from approximately 200 to 1000 nm in radius) (Fig. 6a inset). After switching to global activation, the sizes of these droplets were estimated, and they were tracked for 5 minutes at a frequency of 60 ms. Trajectories were binned based upon size and MSDs, were ensemble averaged, and then fit (Fig. 6a). The subdiffusive exponent *α* was approximately 0.5 for all sizes (Fig. 6b), consistent with a chromatin mesh size below the radius of all interrogated droplets, in agreement with previous estimates^22^. This data also allows us to examine the dependence of the anomalous diffusive coefficient on droplet size (Fig 6c). Interestingly, rather than exhibiting the usual Stokes dependence, *D*_*r*_∼*r*^−1^, the best fit power-law yields *D*_*r*_∼*r*^−0.44±0.18^ with a coefficient of determination of *R*^2^ = 0.45, compared to a *R*^2^ = –0.26 for a power-law slope of −1.

## Discussion

In this study, we sought to understand how the size distribution of intracellular condensates is impacted by their local mechanical environment. We utilized a model system comprised of light-activated condensates, nucleated within the dense viscoelastic chromatin environment in the nucleus of living cells. Consistent with prior results^22^, we found that droplets assembled using intrinsically disordered protein regions (IDRs) from transcriptional proteins tend to localize within chromatin-poor regions, and exclude chromatin as they grow. We found that these model condensates remain relatively stable, even over the span of hours, failing to coarsen into a single large domain, which would be expected for a simple phase-separating system. We find that the viscoelastic chromatin network appears to underlie subdiffusive motion of droplets, which hinders droplet coalescence, giving rise to an anomalously small coarsening exponent.

The two primary mechanisms for droplet coarsening are droplet coalescence (BMC) and Ostwald ripening (LSW), both of which predict a coarsening exponent of *β* = 1/3, significantly larger than our measurement of *β* = 0.12 ± 0.01. We provide evidence that Ostwald ripening does not strongly contribute to the coarsening of droplets. We attribute the lack of Ostwald ripening to a low surface tension, which is expected to be on the order of γ∼*kT*/*ℓ*^2^ where *ℓ* is a characteristic molecular length scale^30^, which for macromolecules (i.e. biomolecules) is relatively large. Indeed, previous work on nucleolar protein condensates in and out of cells suggest that their surface tension is on the order of 10^−7^ N/m^18,31^; considering parameters for FUS_N_ Corelets^27^, we estimate that Ostwald ripening should drive droplet growth at a rate of 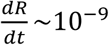 for droplets with an initial radius of 300nm. By contrast, we estimate a significantly larger initial growth rate due to subdiffusive merger: based on an average initial droplet spacing of 1.3 µm (Supplementary Fig. 2), and our tracking data (Fig. 5b), we obtain an approximate growth rate of 6 × 10^−6^µm/s (Supplementary Note). Calculating the average growth rate from our coarsening data (Fig. 1d), we obtain an average growth rate of 2.4 ± 0.3 × 10^−5^ µm/s, within an order of magnitude of this latter, independent estimate.

**Fig. 5:**
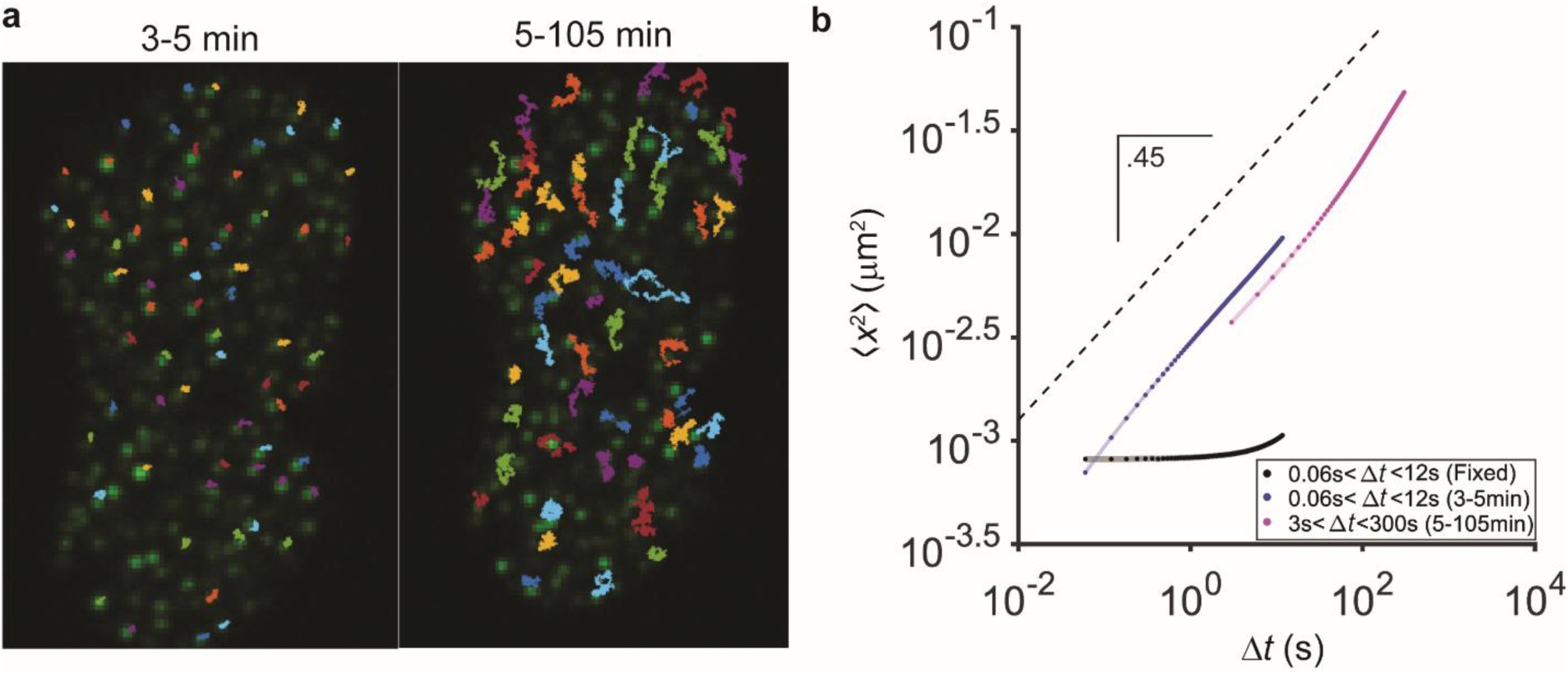
Droplets within the nucleus exhibit strongly subdiffusive motion. **a,** FUS Corelet condensates were tracked at a framerate of 60ms per frame for two minutes (starting 3 minutes after activation). They were then tracked for the following 100 minutes at 3s/frame. Only droplets with continuously visible trajectories were tracked, with a minimum duration of 2 minutes in the short phase and 50 minutes in the long phase. **b,** Mean squared displacements for the short and long tracking phases are shown in blue and magenta respectively, ensemble averaged over 18 cells. Shaded region indicates standard error of the mean. The magenta MSD follows a similar slope but demonstrates nonlinear increase, likely due to stage and whole-nucleus drift. Noise floor (black data points) was estimated to be ∼30*nm* by applying the same tracking protocol on fixed cells, which exhibited similar long-term increases; fitting the MSD of the fixed condensates gave an exponent of .07 which was used as a conservative estimate of the uncertainty for the diffusive exponent.

**Fig. 6:**
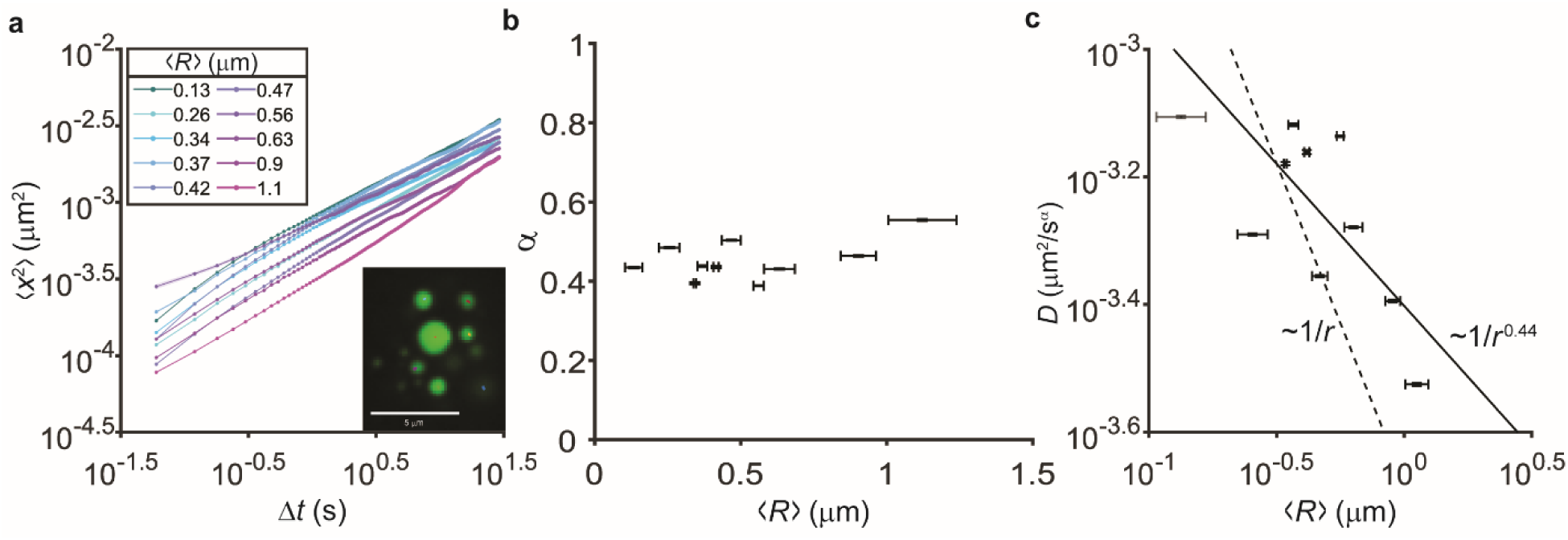
Condensates can be used as microrheological probes of the cellular environment. **a,** Local activation protocols generate size-selectable stable droplets (inset image). After initial stimulation, droplets were imaged for 5 minutes at 60ms/frame and tracked for the entire movie. Trajectories were binned by droplet size and time- and ensembled-averaged to produce an MSD for each size bin. Shaded region indicates standard error of the mean. **b,** Subdiffusive exponent versus droplet size. Horizontal error bar is standard deviation of droplet sizes in each bin; vertical error bar is 95% CI of mean diffusive exponent from power-law fits. **c,** The effective diffusion coefficient decreases with droplet size. Best fit line (solid) is *D*_*r*_∼*r*^-0.44±0.18^ (standard error of the fit; *R*^2^ = 0.45); a *D*_*r*_∼*r*^−1^ reference line is also shown (dashed line; *R*^2^ = –0.26). Horizontal error bar is standard deviation of droplet sizes in each bin; vertical error bar is 95% CI of effective diffusion coefficient from power-law fits. For this fit, the smallest point (⟨*R*⟩ = 0.13 ± 0.3*μm*; plotted in grey) was excluded because its standard deviation overlapped with the diffraction limit significantly.

Ostwald ripening is thus minimal, and instead the coarsening dynamics clearly appear to be dominated by droplet coalescence events, which we reasoned may be retarded by chromatin viscoelasticity. Indeed, from scaling arguments, and our simulation of a fractional Brownian walk, the classic *β* = 1/3 prediction for BMC can be extended to subdiffusive random walks, yielding *β* = α/3, where α is the anomalous diffusion exponent. Consistent with this picture, tracking the motion of our model condensates reveals a high degree of subdiffusive constraint, with α = .48. From this measurement, the *β* = α/3 prediction of *β* ∼0.16, is relatively close to the measured value of *β* = 0.12. The 30% deviation between these numbers could reflect additional physics which we do not account for. For example, the dependence of the diffusive prefactor on droplet size may be non-Stokesian. However, our best fit yields *D*_*r*_∼*r*^−0.5^, which invites a correction of the scaling argument to *β* = α/2.5 (which was confirmed in simulation, Supplementary Fig. 4), increasing the deviation from theory. Instead, we attribute the lower value of the ripening exponent *β* to caging: on very long timescales, we anticipate that the MSD should saturate, due to the presence of boundaries (e.g. heterochromatic domains), further slowing the merger process. This saturation of the MSD may take place on very long timescales which were not captured here due to drift of the nuclei and stage; it is also possible that drift contributed to a small overestimation of the MSD even after correction by image registration. We also note that we tracked condensates and considered them to be entirely independent, while local compressive stresses could cause droplets to slow down when in the vicinity of one another but still merge rapidly when the cavities they induce in the chromatin meet, explaining correlated motions observed in endogenous organelles immediately preceding merger^19,32^.

Interestingly, while the strongly viscoelastic nature of the surrounding chromatin thus appears to retard droplet coalescence, it may also underlie our finding that droplet coalescence is dominant compared to Ostwald ripening. Previous work on synthetic (nonbiological) systems has demonstrated that Ostwald ripening can be suppressed by elastic constraints, stabilizing small droplets even in the presence of large ones^33,34^. Though surface tension drives a preference for the growth of larger droplets, when embedded in a porous elastic matrix, smaller droplets result in smaller deformations and therefore potentially lower energetic costs, dependent on the relative magnitudes of surface tension and matrix elasticity. It has also been shown that the spring constant of chromatin for micron-scale perturbations is ∼10^−3^N/m, four orders of magnitude larger than its surface tension, suggesting that elastic forces could further slow Ostwald ripening in the nucleus.

The intimate link between droplet coarsening dynamics and chromatin viscoelasticity underscores the potential for utilizing these model condensates as passive microrheological probes. Nucleoli and other endogenous condensates have been proposed as probes in a similar fashion^19^, but their dynamics are confounded by essential biological processes, such as nucleation at specific sites and nonequilibrium activity, and by their multicomponent nature. By contrast, our engineered optogenetic droplets can be assembled from IDRs lacking specific interaction domains, and the resulting probe (droplet) size can be controlled using local activation protocols. Within the nucleus, our optogenetic droplets report on the viscoelastic nature of the entangled chromatin matrix. We find strongly subdiffusive dynamics, with α = 0.48 and *D*_*r*_= *r*^−0.44±0.18^, roughly consistent with the Rouse polymer model, which predicts α = 0.5 and *D*_*r*_= *r*^−0.5^. The Rouse model describes the dynamics of a polymer of beads connected by springs and has been used to describe subdiffusion in other biological systems^35^; we speculate that a droplet immersed in a viscoelastic gel and far above the mesh size should exhibit similar dynamics. Future studies will apply this technique to interrogate the mechanical properties of chromatin under varying perturbations.

Our findings also have implications for the assembly of endogenous membraneless organelles. Cajal bodies and nucleoli, associated with relatively chromatin-poor regions, have also been shown to exhibit subdiffusive dynamics, which, moreover, are sensitive to drug treatment, e.g., ATP depletion and transcriptional inhibition^36,37^. Similarly, nuclear speckles display increased mobility as well as increased merger frequency upon transcriptional inhibition and subsequent chromatin condensation. Thus, chromatin condensation and mechanics likely influence the sizes of endogenous membraneless organelles. Future studies will investigate the roles of nonequilibrium biological processes and of specific interactions in the interplay between chromatin mechanics and phase separation.

## Acknowledgements

This work was supported by the NIH 4D Nucleome Program (U01 DA040601, C.P.B.), the Howard Hughes Medical Institute (C.P.B.), and the National Science Foundation, through the Center for the Physics of Biological Function (PHY-1734030) and the Graduate Research Fellowship Program (DCE-1656466, D.S.W.L.). We thank Joshua Riback, Pierre Ronceray, Shunsuke Shimobayashi, Amy Strom, Yaojun Zhang, and the rest of the Brangwynne and Wingreen groups for constructive comments on this work. We also thank Sarah Keller (University of Washington, Department of Chemistry) for useful discussions. We acknowledge Evangelos Gatzogiannis for microscopy assistance and Katherine Rittenbach and the Molecular Biology Flow Cytometry Resource Facility, which is partially supported by the Cancer Institute of New Jersey Cancer Center Support Grant (P30CA072720), for assistance with cell-sorting experiments.

## Methods

### Cell culture

U2OS (a kind gift from Mark Groudine lab, Fred Hutchinson Cancer Research Center) and Lenti-X 293T (Takara) cells were cultured in growth medium consisting of Dulbecco’s modified Eagle’s medium (GIBCO), 10% fetal bovine serum (Atlanta Biologicals), and 10 U/mL Penicillin-Streptomycin (GIBCO), and incubated at 37°C and 5% CO_2_ in a humidified incubator.

### Lentiviral transduction

Lentivirus was produced by transfecting the transfer plasmids, pCMV-dR8.91, and pMD2.G (9:8:1, mass ratio) into LentiX cells grown to approximately 80% confluency in 6-well plates using FuGENE HD Transfection Reagent (Promega) per manufacturer’s protocol. A total of 3 μg plasmid and 9 μL of transfection reagent were delivered into each well. After 60-72 hours, supernatant containing viral particles was harvested and filtered with 0.45 μm filter (Pall Life Sciences). Supernatant was immediately used for transduction or aliquoted and stored at –80°C. U2OS were seeded at 10% confluency in 96-well plates and 20-200 μL of filtered viral supernatant was added to the cells. Media containing virus was replaced with fresh growth medium 24 hr post-infection. Infected cells were imaged no earlier than 72 hr after infection.

### Cell line generation

To generate a population of cells stably expressing H2B-miRFP670, cells were infected in a 6-well plate, transferred to 60mm plate, and finally sorted with a flow cytometer (BD Biosciences), with gating for single-cells intermediate to high levels of miRFP670. The sorted cell line was then infected with viral supernatant as described above to express Corelet components. Gradient activation experiments were performed in this cell line so that the entire nucleus was visible at all times.

### Hoechst Staining

Media was removed from 96 well plate and replaced with fresh media containing 1ug/mL Hoechst 33542 (Thermo) and incubated for 20 minutes prior to imaging.

### Cell fixation

Cells infected with lentivirus and plated on a 96 well plate as described above were placed on an 96 well plate LED array (Amuza) for 15 minutes, after which media was aspirated and 4% paraformaldehyde (Electron Microscopy Sciences, diluted in PBS) was added and the plate remained on the LED array for 15 more minutes. Cells were then imaged and droplets tracked as otherwise described. MSDs were calculated and used to reflect systematic experimental error in the live-cell data.

### Microscopy

All images were taken with a spinning-disk (Yokogawa CSU-X1) confocal microscope with a 100X oil immersion Apo TIRF objective (NA 1.49) and an Andor DU-897 EMCCD camera on a Nikon Eclipse Ti body. A 488nm laser for imaging GFP and global activation, and a 561 was used laser for imaging mCherry. The imaging chamber was maintained at 37°C and 5% CO_2_ (Okolab) with a 96 well plate adaptor. Local activation was performed by using a Mightex Polygon digital micromirror device (DMD) to pattern blue light (488nm) stimulation from a Lumencor SpectraX light engine. All image acquisition was performed using Nikon Elements Advance Research software.

### Image Analysis

#### Image Segmentation

All images were analyzed in Fiji (ImageJ 1.52p)^38^ and MATLAB 2019b (Mathworks). Individual cells were cropped by hand and saved as .tifs, which were analyzed in MATLAB. Briefly, droplets were segmented in the GFP channel by using Otsu’s algorithm to identify the nucleus in the first frame; an intensity threshold was defined as two standard deviations above the mean of the initial nuclear GFP intensity. This threshold was applied to all following frames to identify droplets; regions 4 pixels and smaller were discarded. Then, r*egionprops* was used to identify individual domains and calculate their area. Radii were calculated by dividing the area by π and taking the square root. To calculate volume fraction, the dilute phase of the nucleus was identified by applying a threshold at 3 standard deviations above background concentration (characterized by imaging a sample with no fluorescence). In instances where no initial, pre-activation green channel was available (e.g., local activation experiments), Otsu’s method (*imbinarize* in MATLAB) was used to determine an intensity threshold for droplets. For gradient activation experiments, Otsu’s method was used to segment nuclei in the miRFP channel, and again applied to pixels identified as being inside the nuclei to identify droplets on a per-frame basis. All results were validated by manual inspection. Analysis in Figures 1c, 1d, 2c, 2d, and 5 utilized the same raw data.

#### Particle Tracking and Registration

To track particles image registration was first performed to correct for rigid body motion using the StackReg plugin in Fiji. Subpixel tracking was then performed in Trackmate^39^ using a Laplacian of Gaussians filter-based detector and a blob diameter of 500nm (or an appropriate size for local activation experiments with large droplets), a threshold of 250. Trajectories were then constructed using the simple LAP tracking with max linking and gap-closing distances of 500nm and no frame gap accepted. Only trajectories spanning the entire movie (or in the case of the 100 minute movies, at least half the movie) were accepted. Coordinates were then loaded into MATLAB to calculate MSD and cross-correlations.

### Simulations

Simulations were performed in MATLAB on the Della cluster (Princeton Research Computing). First, 500 droplets of identical radius (1AU) were generated in 3D with periodic boundary conditions. Box size was set based on the volume fraction (generally set at 5%). Overlapping droplets were then merged. Droplet merger was implemented by randomly selecting a pair of overlapping droplets and replacing them with one new droplet centered at their center of mass. The size of the new droplet was size determined by volume conservation of the original pair. This was iterated until no pairs of overlapping droplets remained. Then, for each droplet, a fractional Brownian motion trajectory with the appropriate α was synthesized (*wfbm*) for the appropriate number of timesteps (generally set to 10^4^). The step size was set such that a droplet of radius 1 AU would diffuse an average of 1AU in 3D per timestep. At each timepoint, each droplet proceeded one step along the synthesized trajectory, scaled by the inverse of the droplet size in the case of *D*_*r*_∼1/*r*. Droplets were merged as previously described. Merged droplets ‘inherited’ the predetermined trajectory from one of their parent droplets (chosen arbitrarily), moving with the appropriate step size. This was repeated for the duration of the synthesized trajectories.

## Data Availability

All data from this study are available from the corresponding author upon reasonable request.

## Code Availability

All custom code used in this study are available from the corresponding author upon reasonable request.

