## Supplementary Information for "Chromatin Mechanics Dictates Subdiffusion and Coarsening Dynamics of Embedded Condensates"

### Supplementary Note

*Calculation of coarsening rate by BMC & LSW*

Here, we sought to estimate the growth rate of fully quenched droplets due to both Ostwald ripening and subdiffusive Brownian motion induced coalescence.

*Timescale estimate for Ostwald ripening*

From Lifshitz and Slyozov we can estimate our initial ripening rate as:

$$\frac{dR}{dt} = \frac{2D\gamma c_{sat} v_i^2}{k_B T R^2}$$

Where the surface tension $\gamma \sim 10^{-7}\frac{N}{m}$

Molecular volume $v_i \sim 30nm^3/molecule$ (upper bound)

Saturation concentration $c_{sat} \sim 3uM$

Molecular diffusion coefficient $D = \frac{5um^2}{s}$ (upper bound for “core”; see Bracha *et al*. 2018)

Droplet size $R \sim 0.29um$ (see Supplementary Fig 2);

We therefore find that

$\frac{dR}{dt} \sim 10^{-9} um/s$

*Subdiffusive Brownian Motion-driven Coalescence*

The average merger rate will be given by the spacing divided by the MSD, i.e:

$$D\tau_{merge}^{\alpha} =< (l-2r)^2 >$$

$$\frac{dn}{dt} = -\frac{1}{\tau_{merge}}$$

$$\langle\frac{dn}{dt}\rangle = -\left(\frac{D}{(< l(t) > -2 < r(t) >)^2}\right)^{1/\alpha} \sim 2 \times 10^{-5}/s$$

$$\langle l(t_0)\rangle \sim 1.3um$$

$$\langle R(t_0)\rangle \sim 0.29um$$

$$\langle x^2(\tau)\rangle \sim 10^{-2.5} \mu m^2/s^{0.47}$$

$$\langle\frac{dR}{dt}\rangle \sim < \mathrm{R} > \langle\frac{dn}{dt}\rangle$$

$$\langle\frac{dV}{dt}\rangle \sim 6 \times 10^{-6}\ \mu m/s$$

From our data, we observe a change from $10^{-.35}$ to $10^{-.45}$ $\mu m$ over 102 minutes, giving an average growth rate of approximately $10^{-5}$ $\mu m$/s, within an order of magnitude of our independently calculated estimate of merger-based growth rate from tracking data.

**Supplementary Figure 1.** Analysis from Figs. 2 and 3c were repeated using an integrated intensity metric to estimate the size of droplets. **a,** Average integrated droplet intensity grows as a power law in time. **b,** Nondimensionalizing with $t_0 = 3$min, averaging, and fitting gives an exponent of 0.12. **c,** Integrated intensity is conserved among collisions. **d,** Mean squared error calculated by assuming that volume must be conserved among collisions and then determining the deviation of the final volume post collision from the prediction. The root mean square error is similar but slightly greater for the integrated intensity method.

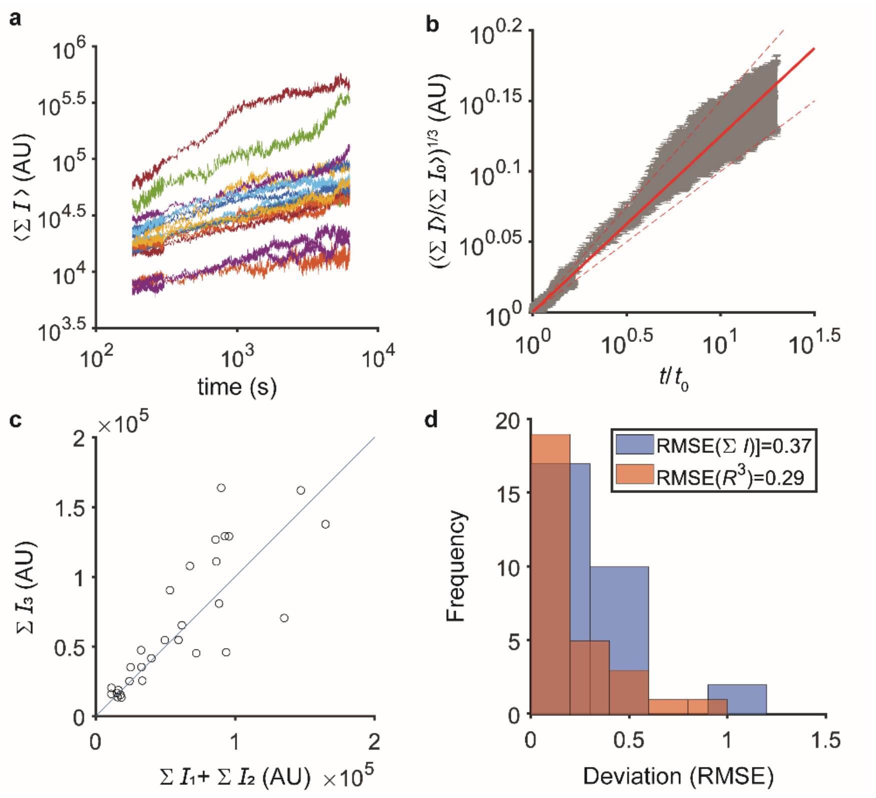

**Supplementary Figure 2.** We characterized our system at early times immediately following a blue-light quench by taking high-speed images for the first five minutes of activation. **a,** First, supersaturation was calculated by estimating the intensity in the nucleus but outside droplets and dividing by the average nuclear intensity in the first frame (i.e., before any visible condensation). This was averaged over cells for each timepoint; shading represents standard error of the mean. Supersaturation was by 180 seconds. **b,** To quantify nucleation dynamics number of droplets was calculated at each frame following activation; a quick increase and then saturation and slow decrease is observed as droplets begin to coalesce. For each cell, the number of droplets was taken at each frame, divided by the time average for that cell, and then averaged over time. Error bars reflect standard error of the mean across cells. **c,** To characterize the heterogeneity of coarsening behavior across cells, we individually fit the average droplet radius versus time for each cell. We computed the volume fraction of droplets by dividing of the number of pixels containing droplets at the final frame of activation (105 minutes) to the total number of pixels in the nucleus in the first frame of activation. This volume fraction was plotted against the coarsening rate fit for individual cells (error bar is 95% CI); the noise in the coarsening behavior decreases with volume fraction to an average value of 0.12. **d,** Finally, to estimate the average rate of growth due to subdiffusive merger, we compare minimum inter-droplet distance per droplet versus average radius of droplets at $t_0$ per cell and find that due to existing in a similar volume fraction regime and having a similar nuclear size, we observe a strong correlation. Error bars reflect SEM.

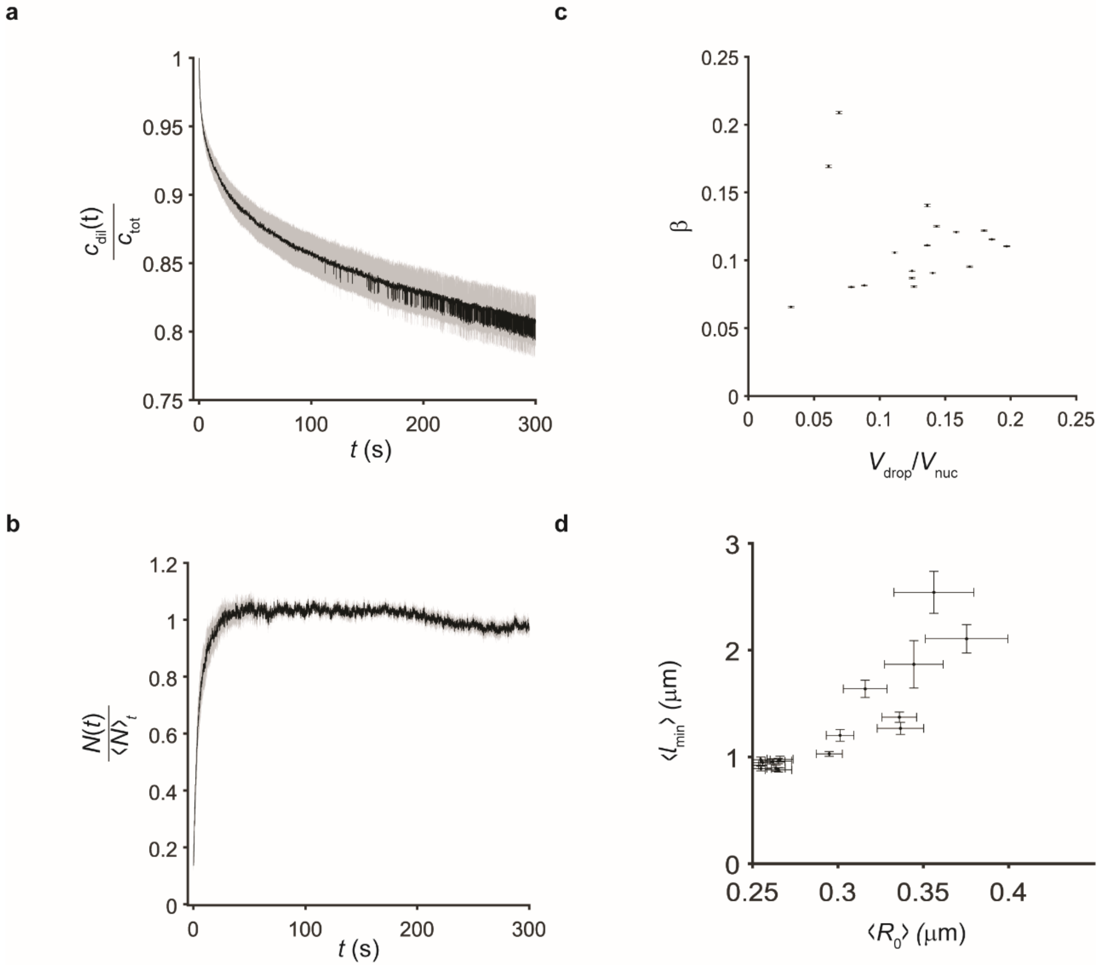

**Supplementary Figure 3.** Equilibration of saturated concentration in Figure 4d,e is fast. Nuclei from Figure 4D were segmented and pixel intensity in the dilute phase was binned by location with respect to the gradient of activation, normalized, averaged over cells, and plotted. Cells were activated using a blue light intensity gradient for 30 seconds; at the end of that time, a gradient is observed in the dilute concentration favoring the side of the nucleus experiencing higher intensity stimulation (black line). Within one frame (3 seconds) following the switch to global activation concentration of protein in the dilute phase becomes uniform (blue line). By 6 seconds(dark red line), small sub-diffraction-size droplets begin to nucleate in the previously unactivated half of the nucleus, accounting for the slight increase in intensity which persists in the 9 second profile (red line).

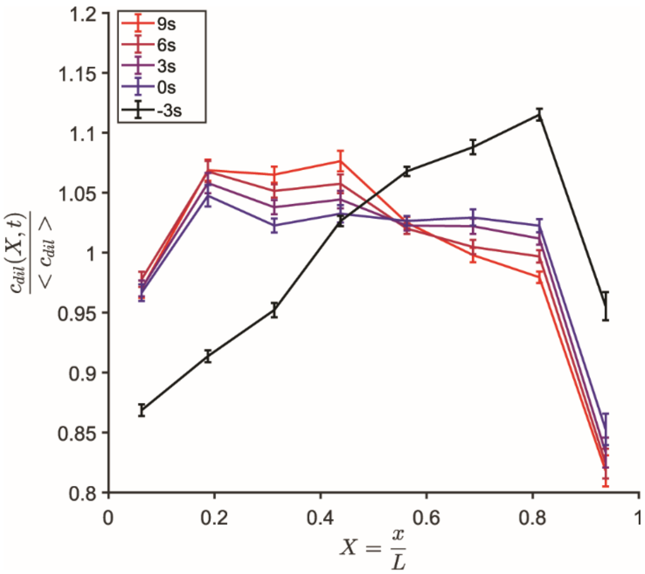

**Supplementary Figure 4.** Scaling behavior simulated with $D \sim r^{-.5}$. **a,** As in Figure 4, values of *α* were chosen ranging from 0.1 to 1 with 20 replicates per condition, and merged over $10^4$ timesteps. Step size was scaled as $r^{-.5}$. Average radius of droplets in each replicate was averaged, for each condition over 20 replicates and plotted. Shaded error bar reflects standard error of the mean. Power laws were fit between $10^2$ and $5 \times 10^3$ timesteps. **b,** The input α for each condition was plotted against the calculated β and fit with a line with 0 y-intercept, revealing a proportionality of 0.40 according to the fit.

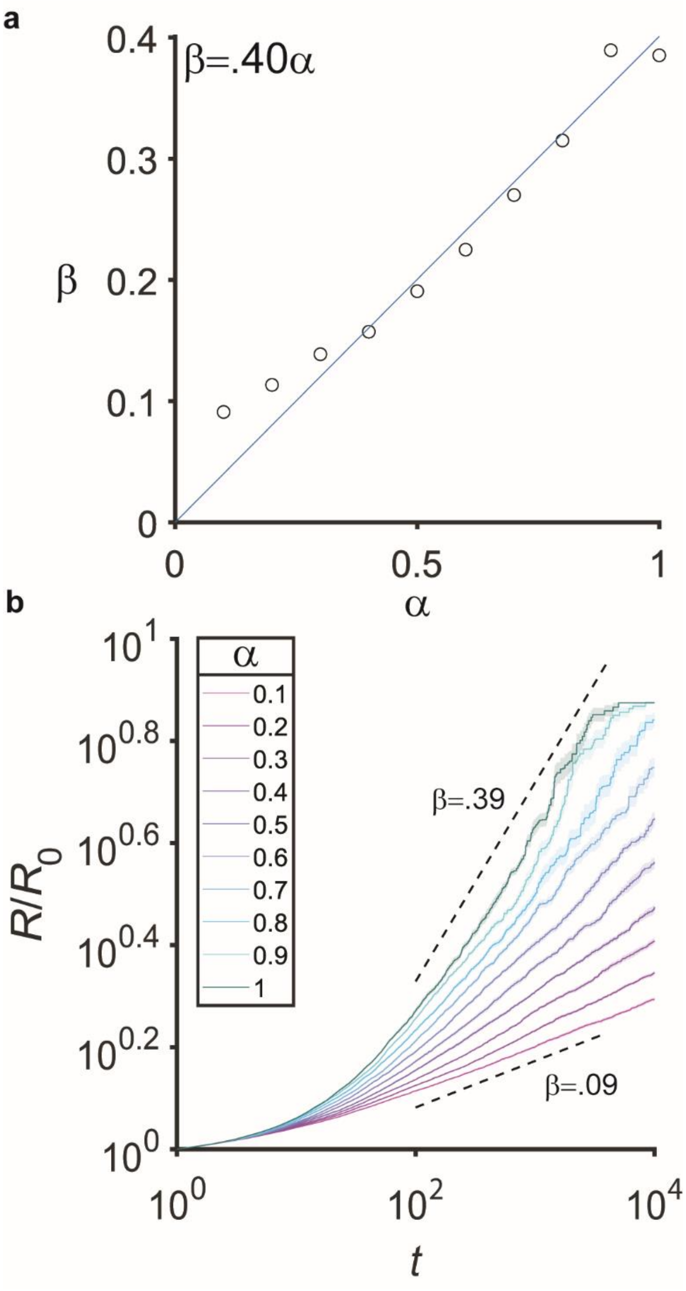
